# An accessible transfection protocol for choanoflagellates

**DOI:** 10.64898/2026.03.10.710884

**Authors:** Maria H.T. Nguyen, Iliana S. Hernandez, Florentine U. Rutaganira

## Abstract

Choanoflagellate genetics has undergone rapid and impactful developments in the last decade. Currently, the primary method for genetic modification of choanoflagellates relies on proprietary nucleofection reagents to deliver transgenes for ectopic expression or CRISPR-Cas9 ribonucleoprotein complexes for targeted genome editing. The acquisition of proprietary buffers required for nucleofection can hamper advances in choanoflagellate research due to costs, shipping limitations, and restrictions that prevent buffer components from being optimized for understudied organisms. Therefore, we test whether a low-cost in-house electroporation buffer developed for other systems can replace the proprietary buffer currently used for choanoflagellate transfection. Here, we present an in-house buffer with transfection efficiency comparable to that of the previously established proprietary buffer. This work increases the accessibility of choanoflagellate genetics and can broaden research participation in investigating animal origins.

## Introduction

Choanoflagellates are the closest living relatives of animals, and are therefore phylogenetically positioned to be a powerful model system for investigating the origins of animal multicellularity and development. Understanding the molecular history of how animal body plans and cell types first evolved requires tools that enable precise genetic manipulation of choanoflagellates, and in the last decade, we have seen impactful advances toward this goal.

Early genetic manipulation of choanoflagellates was achieved through forward genetic screens, which enabled the identification of genes controlling the multicellular development of the choanoflagellate *Salpingoeca rosetta*^1^. A few years later, a method using the Lonza 4D Nucleofector to enable stable transfection and transgene expression was established, thereby opening the door to reverse genetics^2^. This work was further advanced through CRISPR-Cas9-based genome editing in choanoflagellates by delivering a CRISPR-Cas9 ribonucleoprotein complex, along with an oligonucleotide repair template, to direct precise DNA cleavage and homologous recombination at a target site^3^. Successfully edited cells were enriched through cycloheximide selection by co-editing *rpl36a* to confer resistance^3^. More recently, an approach that links gene targeting and selection to the same locus increased selection efficiency for gene knockouts by coupling homologous recombination of a selectable marker flanking translation termination sequences at the target site^4^. Notably, this modified approach demonstrated that repair templates for inserting the selectable marker and translation termination sequences can be generated via PCR, thereby eliminating the need for commercially synthesized oligonucleotides and offering a more cost-effective knockout strategy.

While these advances have greatly expanded the capacity to study choanoflagellate gene function through knockout and overexpression, a key limitation remains: the nucleofection (an electroporation-based transfection method) kits required for these protocols rely on proprietary buffers, which are expensive and can restrict research accessibility. For instance, the costs associated with routine nucleofection may hamper large-scale experiments or broader adoption of choanoflagellate genetics. Second, nucleofection buffers were developed specifically for mammalian cell lines; therefore, the use of proprietary buffers for nucleofection prevents systematic modification of buffer components to be optimized for understudied organisms that may have different transfection requirements. Building on prior work by choanoflagellate researchers and transfection approaches that use in-house electroporation buffers developed in other model organisms, we sought to further optimize the transfection protocol for cost-efficiency and to identify an in-house electroporation buffer with transfection efficiency comparable to that of the previously established method using a proprietary nucleofection buffer. As previously established^2^, we used nanoluciferase expression as a readout of transfection efficiency and optimized pulses for promising in-house electroporation buffers. We identify several candidate buffers and demonstrate that, with further optimization, the in-house electroporation buffer achieves transfection efficiencies comparable to the proprietary standard. This work lowers the barrier to entry for choanoflagellate genetics and increases accessibility for laboratories investigating animal origins.

## Methods

### Choanoflagellate culturing

*Salpingoeca rosetta* was maintained in monoxenic co-culture with the bacterium *Echinocola pacifica* (co-culture “SrEpac”; American Type Culture Collection [ATCC], Manassas, VA; Cat No. PRA-390). Cells were grown in High Nutrient Media, which contained 4% (v/v) Artificial Known SeaWater Complete (AKSWC) and 4% (v/v) Artificial Known Cereal Grass Media 3 (AKCGM3) in Artificial Known SeaWater (AKSW)^3^.

### Preparation of Chicabuffers

Each Chicabuffer buffer component was weighed and stored in separate 15 mL conical tubes at room temperature. During papain treatment of choanoflagellate cells, solutions of the individual components of the selected Chicabuffer^5,6^ were prepared freshly in milliQ water as 10X concentrated stocks in 10 mL volumes (Supplementary Table S5) and 0.2 µm filter-sterilized. The respective salts, additives, and phosphate buffer were then assembled in a microcentrifuge tube and kept on ice.

### SpCas9 Ribonucleoprotein (RNP)

CRISPR RNA targeting *rpl36a* and oligonucleotides used as repair templates^3^ to render cells cycloheximide-resistant were ordered from IDT. Upon arrival, guide RNAs and repair oligonucleotides were prepared as previously described^3^. Briefly, gRNAs were prepared by annealing crRNA (IDT, CTTCTTGCGGAAGATGGGCT) with tracrRNA (IDT, Cat No. 1072532). Lyophilized oligonucleotides (GCACGGAACATCGCCTCCAATGACATGCATGAGACAAGCACCTTCTTGCGGAAGATctgCTTGGTCTGACCACCGAAACCGCTCTGCTTCCGGTCGTAACGACGCTTACCTACACACACG) were resuspended to a concentration of 250 µM in 10 mM HEPES-KOH, pH 7.5, and incubated at 55°C for 1 hr. The gRNA and repair oligonucleotides were stored at −20°C. On the day of nucleofection, SpCas9 RNP was prepared by incubating 2 µL of SpCas9 (NEB, Cat No. M0646M) with 2 µL of crRNA at room temperature for 1 hr.

### Transfection Protocol

48 hours before electroporation, a 200 mL SrEpac culture was seeded at 8,000 cells/mL in High Nutrient Media in T-300 Suspension Culture Flasks with Vent Cap (Celltreat Scientific, Cat No. 229540), and incubated at 22°C with 60% humidity.

1. *Choanoflagellate wash* On the day of nucleofection, the full volume of the SrEpac culture was harvested, split into 4 50 mL conical tubes, and spun at 2,400 × g for 5 minutes at 4°C. The supernatant was discarded and the pellet was resuspended in 50 mL AKSW. The wash steps were repeated for a total of 3 AKSW washes. For the final wash, the supernatant was carefully removed to maximize bacterial clearance, then the pellet was resuspended in 100 µL AKSW. Cell concentration was quantified by diluting 2 µL of washed cells into 196 µL AKSW, then adding 2 µL of 16% formaldehyde (Thermo Scientific™, Cat No. PI28908), and counting 10 µL of the mixture using a LUNA-II™ automated cell counter (Logos Biosystems). Cells were diluted to 5E7 cells/mL and split into 100 µL aliquots.
2. *Glycocalyx digestion* The aliquots were spun at 800 × g for 5 minutes at 4°C. During this time, the priming solution was prepared: 5 µL of papain (MilliporeSigma, Cat No. P3125) was added to the dilution buffer (50 mM HEPES-KOH, pH 7.5, 200 mM NaCl, 20% (v/v) glycerol, 10 mM L-cysteine; 0.2 µm filter-sterilized and stored at −70°C). Then, 6 µL of the dilution buffer with papain was added to 400 µL of priming buffer (40 mM HEPES-KOH, pH 7.5, 34 mM lithium citrate, 50 mM L-cysteine, 15% PEG 8000; 0.2 µm filter-sterilized and stored at −70°C) and kept on ice. The cells were immediately removed from the centrifuge, resuspended in 100 µL of priming solution, and incubated at room temperature for 35 minutes. The digestion was quenched by adding 10 μL of 50 mg/mL bovine serum albumin (dilution buffer without L-cysteine; GoldBio, Cat No. A-420-10). Then, cells were harvested by centrifugation at 1,250 × g for 5 minutes at 4°C and resuspended in 25 µL of electroporation buffer. The resulting suspension is referred to as “primed cells”.
3. *Electroporation* CRISPR-Cas9 electroporation reactions were prepared in microcentrifuge tubes or PCR strip tubes, which consisted of: 4 μL SpCas9/gRNA RNP, 2 μL oligonucleotide repair template, 2 μL primed cells, and 2 µL of 20 mg/mL pUC19 plasmid (gift from the Nicole King Lab, UC Berkeley), brought to a final volume of 24 μL with nucleofection buffer. Plasmid electroporation reactions were prepared in microcentrifuge tubes or PCR strip tubes, which consisted of: 1 µL of nanoluciferase plasmid, 2 μL primed cells, 2 µL of 20 mg/mL pUC19 plasmid (gift from the Nicole King Lab, UC Berkeley), 1 µL of 100 mg/mL Heparin (in milliQ H_2_O, SigmaAldrich, Cat No. H3149-100KU), and ATP (0.2 µm filter-sterilized in milliQ H_2_O, GoldBio, Cat No. A-081-25), brought to a final volume of 23 µL with nucleofection buffer. The electroporation mix was transferred into nucleocuvette wells and pulsed using the Lonza 4D Nucleofector X unit. Immediately after pulsing, 100 µL of ice-cold recovery buffer was added to each reaction and mixed by gently tapping the cuvette. After a 5-minute recovery, the entire contents of each nucleocuvette were transferred into either 6-well plates (CRISPR-Cas9) or 12-well plates (plasmid transfection) containing 2 mL or 1 mL, respectively, of Low Nutrient Media, which contained 1.5% (v/v) AKSWC and 1.5% (v/v) AKCGM3 in AKSW. The cells were incubated at 22°C and 60% humidity for 30 minutes, then fed 5 µL of 10 mg/mL *E. pacifica* and incubated for another 24 hours before downstream experiments.

### Cycloheximide selection

After allowing *S. rosetta* electroporated with SpCas9 RNPs and repair oligonucleotides for *rpl36a*^*P56Q*^ to recover for one day, 10 µl of 1 µg/ml cycloheximide was added to the 2 mL culture of transfected cells. The cells were incubated with cycloheximide for 5 days prior to observation.

### Nanoluciferase plasmid

Pulse optimization was performed using pNK809, a plasmid encoding *S. rosetta* codon-optimized Firefly and NanoLuc luciferases under a strong *EFL* promoter (gift from the Nicole King Lab, UC Berkeley, Addgene plasmid # 196406)^7^. The plasmid was prepared by Genscript (Plasmid DNA Preparation Service) at 5 mg/mL in nuclease-free water and stored at −20°C.

### Nanoluc reporter assay

Transfection efficiency was assessed by measuring nanoluciferase luminescence using the Nano-Glo Luciferase Assay System (Promega, Cat No. N1110). The assay buffer was thawed at room temperature. Cells were collected from each well of the transfection plate by pipetting up and down 2–3 times with a P1000 and transferring to individual microcentrifuge tubes. Samples were centrifuged at 4,200 × g for 10 minutes at 4°C, after which the supernatant was removed and each pellet was resuspended in 50 µL of Nano-Glo lysis buffer. The Nano-Glo working solution was prepared by combining the substrate with assay buffer at a 1:25 ratio. Each resuspended sample was gently vortexed, then 12.5 µL was transferred to a well of a 384-well white flat-bottom plate (Greiner, Cat No. 781080) containing 12.5 µL of Nano-Glo working solution, and the mixture was mixed by gentle pipetting to prevent bubbles. Luminescence was read immediately on a plate reader (PerkinElmer (now Revvity) Victor Nivo Alpha S) with the plate lid removed. Data were normalized to the average of no-pulse controls, with blank subtraction applied where no-pulse control signals exceeded background.

### Optimization of Pulses

Pulse programs were selected using the Lonza fine-tuning matrix (https://bioscience.lonza.com/download/content/asset/31152; accessed 6 March 2026). Beginning with the suggested ‘best program,’ pulse conditions were varied along axes of increasing viability and increasing efficiency to identify the program that maximized transfection efficiency.

### Reusing of Cuvettes

To reduce waste and costs associated with nucleofection even further, we reuse our nucleocuvettes, taking inspiration from (https://dx.doi.org/10.17504/protocols.io.64mhgu6) with the following modifications: during the 30-minute post-nucleofection recovery period, wells were rinsed with 70% ethanol, incubated for 5 minutes, and emptied by inversion and vigorous shaking, followed by a milliQ H_2_O rinse. This ethanol-and-water cycle was repeated for a total of 3 rinse cycles to maximize the removal of residual salts and cellular debris that could interfere with subsequent nucleofections. We stored cuvettes overnight in >95% ethanol, removed them the next day, and let them dry before long-term storage in a covered vessel to prevent dust build-up. On the day of nucleofection, we repeat the 70% ethanol and milliQ water rinse steps, with the final rinse performed under a sterile hood. The nucleocuvettes were allowed to air dry completely before use.

## Results

### Preliminary In-House Electroporation Buffer Testing

Since prior work demonstrated that the percentage of genome editing correlates closely with transfection efficiency^3^, we first sought to determine whether Chicabuffers^5,6^ could functionally replace the proprietary Lonza SF buffer using a pass/fail assay before pursuing quantitative optimization. We performed transfection with CRISPR-Cas9 RNP complexes as previously described^3^ using various in-house electroporation buffers and selected for successfully edited cells (*rpl36a*^*P56Q*^) by culturing the resulting transfected *S. rosetta* in the presence of cycloheximide for five days. Cells were observed for evidence of survival and proliferation as an indicator of successful genome editing (Figure 1A). Across five trials, we identified two in-house electroporation buffer candidates: Chicabuffer 1M and 3H, which supported successful editing in at least two of five trials (Figure 1B). These results established 1M and 3H as viable candidates for further testing as replacements for the Lonza SF buffer.

**Figure 1.**
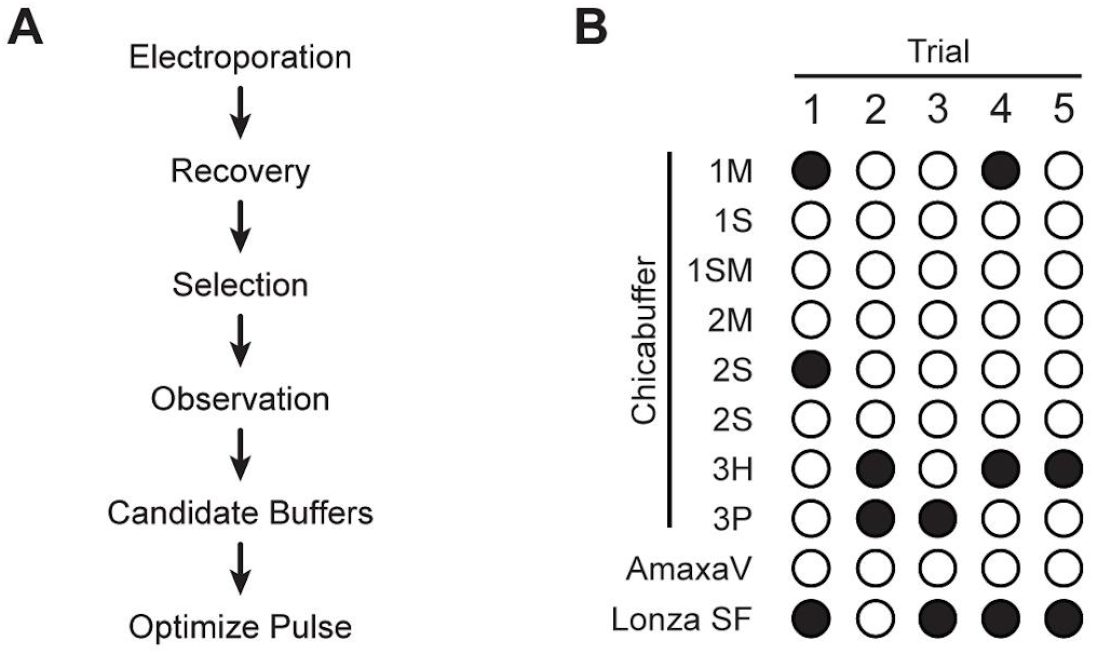
Preliminary trials identify Chicabuffers 1M and 3H as candidates in-house electroporation buffers for further optimization. (**A**) Schematic overview of the post-electroporation workflow used to identify candidate buffers. (**B**) Chicabuffers and AmaxaV buffers were evaluated across five independent trials, with three transfection reactions per buffer. Filled circles indicate *Salpingoeca rosetta* survival in one of three transfections after 5 days of antibiotic selection.

### Narrowing Candidates between 1M and 3H

To quantitatively compare the transfection efficiency of the two candidate buffers, we used the Lonza nucleofection optimization program matrix and assessed efficiency using a luciferase (NanoLuc) reporter assay across the ‘best programs’ in the matrix (Figure 2A-B)^2^. 1M consistently outperformed 3H in transfection efficiency, establishing 1M as the preferred candidate for further optimization.

**Figure 2.**
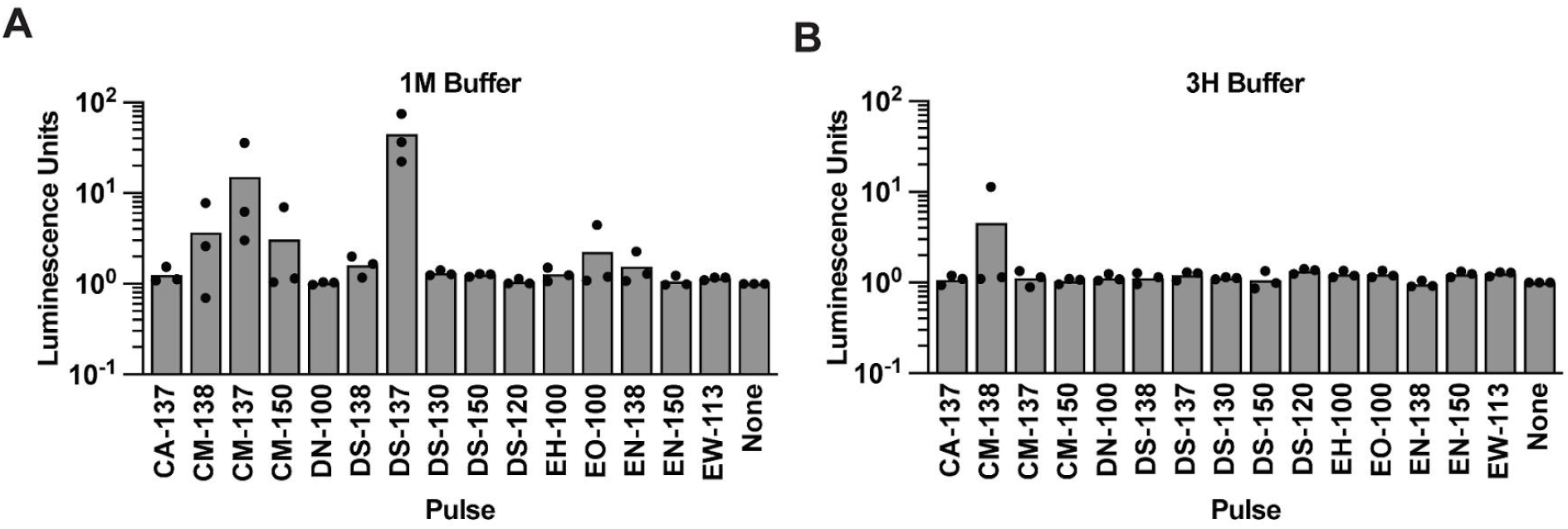
Chicabuffer 1M with pulse DS-137 yields the optimal combination of in-house buffer and pulse program for further fine-tuning of transfection efficiency. Plasmid-encoded nanoluciferase was transfected into *S. rosetta* using (**A**) Chicabuffer 1M and (**B**) Chicabuffer 3H across a range of pulse programs varying in efficiency and viability in triplicate. Nanoluciferase activity was measured 24 hours post-nucleofection and reported as luminescence relative to the average of three no-pulse controls.

### Identification of the Ideal Buffer and Pulse Conditions

To maximize the performance of 1M, we further optimized the nucleofection pulse program by fine-tuning conditions to improve both cell viability and transfection efficiency. This analysis identified pulse DG-137 as the optimal program for use with 1M (Figure 3A). Using their respective optimized pulse conditions, we directly compared 1M to the proprietary Lonza SF buffer in a head-to-head transfection and found comparable transfection efficiencies between the two buffers (Figure 3B). Additionally, we were curious whether fully assembled 1M or its stock components could be stored frozen and thawed prior to use. Our results indicate that freshly prepared 1M yields higher transfection efficiency (Figure S1). This could be due to freezing altering solubility. Together, these results demonstrate that 1M can serve as a cost-effective alternative to the Lonza SF buffer for nucleofection of *S. rosetta*.

**Figure 3.**
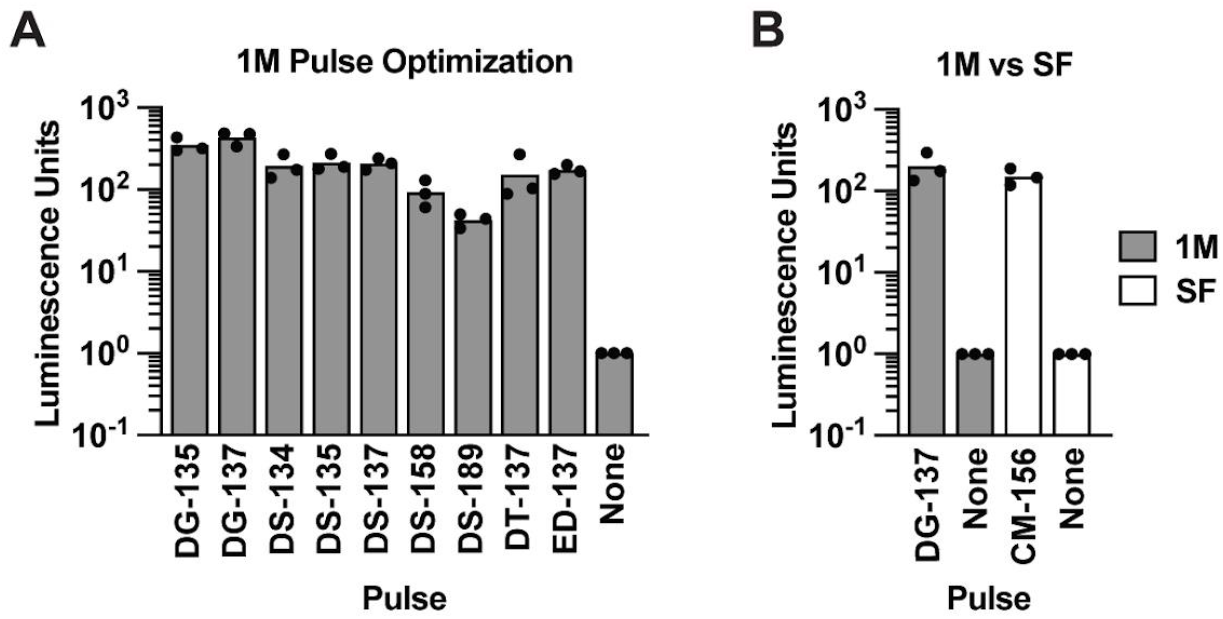
Chicabuffer 1M with pulse DG-137 achieves transfection efficiency comparable to the previously established Lonza SF buffer and CM-156 pulse combination. (**A**) Transfection pulse was fine-tuned by testing alternative pulse programs with greater viability (pulses left of DS-137) or greater efficiency (pulses right of DS-137), as measured by nanoluciferase activity. As in Figure 2, transfection was performed in triplicate. (**B**) Nanoluciferase activity comparing Chicabuffer 1M with optimized pulse DG-137 against the proprietary Lonza SF buffer with pulse CM-156.

## Discussion

Here, we demonstrate that the in-house electroporation Chicabuffer 1M can achieve transfection efficiencies comparable to the proprietary Lonza SF buffer when paired with an optimized pulse program, providing a cost-effective alternative for choanoflagellate genetic manipulation.

One important caveat to our initial screening approach for the preliminary buffer comparisons is the use of pulse CM-156 during electroporation, a program previously optimized for the Lonza SF buffer rather than for each individual in-house electroporation buffer. As a result, our screen to detect *S. rosetta* growth in cycloheximide after CRISPR-Cas9 editing of *rpl36a* may not have captured the full potential of all in-home electroporation buffers tested. It is possible that additional candidates, including those that did not meet our threshold in early trials, could achieve comparable or superior efficiencies if subjected to independent pulse optimization. Furthermore, changes in culturing conditions (e.g., nutrient level, temperature) alter choanoflagellate growth and will also require optimization of transfection conditions.

Therefore, we recommend that researchers who alter culturing conditions or seek to optimize transfection efficiency in another organism perform a full-factorial optimization of electroporation buffer composition and pulse conditions. Additionally, those interested in batch-preparing in-house buffers for future work should also test the transfection efficiency of fully assembled electroporation buffers or their stock components stored at room temperature or 4°C, as suggested by Lonza for their nucleofection buffers, including SF.

A remaining limitation of the current protocol is the continued requirement of the Lonza 4D Nucleofector and nucleofection kit (nucleocuvettes are not sold separately), which remains a barrier for some laboratories. One promising avenue to further increase accessibility is to explore the two-pulse electroporation method^8,9^, which may enable transfection with more widely available electroporation equipment and further reduce the cost of entry for transfection in non-traditional model systems.

## Author Contributions

M.H.T.N. and F.U.R designed experiments; M.H.T.N. and I.S.H. performed nucleofections; M.H.T.N. performed nanoluciferase assays; M.H.T.N. and F.U.R. analyzed data; and M.H.T.N. and F.U.R. wrote the manuscript.

## Data Availability

All raw data are included in the supplementary materials (Table S2-5) for Figures 2–3 and Figure S1.

## Acknowledgements

We thank Jesus Espinoza from the Booth Lab and David Booth for nucleofection training. We also thank the following choanoflagellate researchers for generously answering questions and giving advice about the nucleofection protocol, particularly during troubleshooting: Thibaut Brunet and lab, Maxwell Coyle, Alain Garcia De Las Bayonas, Michael Carver, and Jeffrey Colgren. We thank the King Lab (UC Berkeley, CA) for pUC19 and pNK809, the Lingyin Li Lab (Stanford University School of Medicine, CA) for the Lonza 4D Nucleofector. M.H.T.N. is supported by the Howard Hughes Medical Institute (HHMI) Gilliam Fellowship, Ford Foundation Predoctoral Fellowship, National Science Foundation Graduate Research Fellowship, and ChEM-H Chemistry-Biology Interface Fellowship. F.U.R. is a Biohub, San Francisco Investigator and is also supported by the Howard Hughes Medical Institute (HHMI) Hanna H. Gray Fellowship. M.H.T.N. extends special gratitude to Ernesto Bonilla and Bear for their unwavering support.

---

**Supplementary Figure 1.**
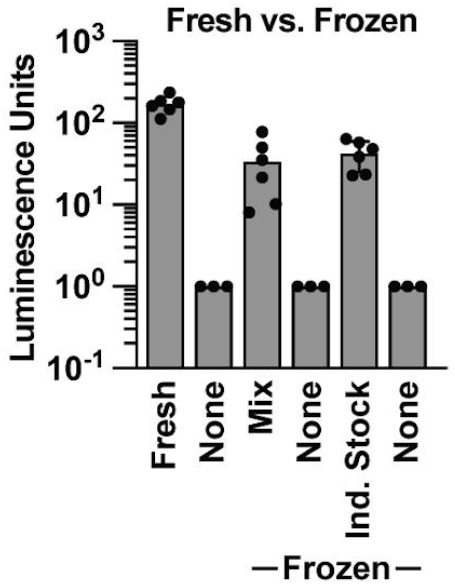
Transfection efficiency was compared across freshly prepared, frozen, and Chicabuffer 1M assembled directly from frozen individual stock components using pulse DG-137. Freshly prepared Chicabuffer 1M shows higher transfection efficiency compared to frozen buffer or frozen individual stock components.

